# Mechanisms and neuroanatomy of response selection in tool and non-tool action tasks: Evidence from left-hemisphere stroke

**DOI:** 10.1101/2022.11.30.518581

**Authors:** Frank E. Garcea, Laurel J. Buxbaum

## Abstract

The ability to select between potential actions is central to the complex process of tool use. After left hemisphere stroke, individuals with limb apraxia make more hand action errors when gesturing the use of tools with conflicting hand actions for grasping-to-move and use (e.g., screwdriver) relative to tools that are grasped-to-move and used with the same hand action (e.g., hammer). Prior research indicates that this grasp-use interference effect is driven by abnormalities in the competitive action selection process. The goal of this project was to determine whether common mechanisms and neural substrates support the competitive selection of task-appropriate responses in both tool and non-tool domains. If so, the grasp-use interference effect in a tool use gesturing task should be correlated with response interference effects in the classic Eriksen flanker and Simon tasks, and at least partly overlapping neural regions should subserve the 3 tasks. Sixty-four left hemisphere stroke survivors (33 with apraxia) participated in the tool- and non-tool interference tasks and underwent T1 anatomical MRI. There were robust grasp-use interference effects (grasp-use conflict test) and response interference effects (Eriksen flanker and Simon tasks), but these effects were not correlated. Lesion-symptom mapping analyses showed that lesions to the left inferior parietal lobule, ventral premotor cortex, and insula were associated with grasp-use interference. Lesions to the left inferior parietal lobule, postcentral gyrus, insula, caudate, and putamen were associated with response interference in the Eriksen flanker task. Lesions to the left caudate and putamen were also associated with response interference in the Simon task. Our results suggest that the selection of hand posture for tool use is mediated by distinct cognitive mechanisms and partly distinct neuroanatomic substrates from those mapping a stimulus to an appropriate motor response in non-tool domains.

## Introduction

A central challenge for the brain is to select a specific action from among a range of possible actions for motor execution (Cisek & Kalaska, 2010; Gold & Shadlen, 2007; Kim et al., 2021; Rushworth et al., 2009). Prior research suggests that the sensory-motor system may prepare multiple possible actions in parallel while accumulating evidence to bias processing toward appropriate goal-directed actions (Cisek, 2007; Cisek & Kalaska, 2005; Gallivan et al., 2016; Kim & Shadlen, 1999; Song, 2017). Selecting among candidate actions is resource-demanding, and typically exerts a processing cost in terms of accuracy and/or response time.

For humans, the ability to skillfully manipulate tools across contexts and environments presents a unique challenge for the action selection system: tool use requires selection of a specific hand posture (or set of postures), and these may be distinct from the hand posture(s) appropriate for grasping a tool to move it. Prior studies have shown that in response to images of tools, representations of tool shape and size—visuomotor properties relevant for tool-directed grasping to move the tool—are evoked more rapidly than representations subserving functional tool manipulation (Lee et al., 2013). In neurotypical individuals, these differences in activation time-course can interfere with the production of tool use actions. For example, participants are slower to initiate a tool-use action to tools with incongruent hand postures for grasping and use (e.g., screwdriver) relative to tools with a congruent hand posture for grasping and use (e.g., hammer) (Jax & Buxbaum, 2010; Osiurak et al., 2013). Such effects of grasp-use interference disappear when tools are placed beyond reach, suggesting that the observed processing cost is attributable to activation of competing motor plans for interacting with tools (Kalenine et al., 2016; see also Wamain et al., 2018).

Individuals who exhibit limb apraxia (hereafter, apraxia) after left hemisphere stroke are impaired when asked to pantomime tool use despite relatively intact tool-directed reaching and grasping-to-move (Buxbaum, Johnson-Frey, et al., 2005; Randerath et al., 2010; Rounis & Humphreys, 2015). Prior studies have demonstrated weak and delayed activation of tool use representations in apraxia (Garcea et al., 2019; Lee et al., 2014; Myung et al., 2010). These reduced and delayed activations render apraxic participants susceptible to abnormal grasp-on-use interference and increased hand posture errors when they pantomime the use of tools with incongruent hand postures for grasping and use relative to tools with congruent hand postures for grasping and use (Jax & Buxbaum, 2013).

The relation between susceptibility to grasp-on-use interference in tool tasks and response competition in non-tool response competition tasks has not, to our knowledge, been investigated. Our first objective was to assess the relation between the tool grasp-use interference effect and response-interference effects in the well-studied Eriksen flanker (Eriksen & Eriksen, 1974) and Simon tasks (Simon, 1969) in 64 left cerebrovascular accident (LCVA) participants (33 of whom exhibited apraxia). The Eriksen flanker task (hereafter, flanker task), requires participants to categorize a central stimulus by means of a motor response (button press) while inhibiting a motor response invoked by flanking stimuli on each side. Response interference is greater when the flanking stimuli are incongruent with the central stimulus, resulting in longer response times relative to a congruent condition.

In one version of the Simon task, an arrow is presented on the left or right of the display and participants indicate whether the arrow is facing left or right by means of a left or right button press, respectively. There is greater response interference when the motor response is incongruent with the side on which the arrow is presented (e.g., left button press when the arrow is presented on the right, and vice versa), resulting in longer response times relative to a congruent condition. We investigated whether we would observe correlations between interference effects across tool and non-tool tasks, as has been shown in some studies of flanker and Simon tasks with neurotypical individuals (e.g., Whitehead et al., 2019; but see Hedge et al., 2021; Rey-Mermet et al., 2018; Wager et al., 2005). If so, this would provide evidence of a common mechanism across tool and non-tool action domains enabling the selection of a task-appropriate motor responses in the face of competition.

Our second objective was to assess the relation between brain regions critical for tool-related response selection and the brain regions associated with flanker and Simon response interference effects. In prior studies, the lesion sites associated with the grasp-use interference effect have included the left inferior parietal lobule, and in particular, the supramarginal gyrus (Watson & Buxbaum, 2015), the left inferior frontal gyrus bordering on the left ventral premotor cortex (Garcea et al., 2019), and the left insula (Dressing et al., 2021; Watson et al., 2019). These regions are nodes in the ventro-dorsal stream of the left hemisphere (the “Use” system), as distinct from the bilateral superior parietal, intraparietal sulcus, and dorsal premotor regions that comprise the dorso-dorsal stream (the “Move” system) (Binkofski & Buxbaum, 2013; Rizzolatti & Matelli, 2003). We have previously suggested that the left inferior parietal lobule maintains grasp and use hand posture representations in working memory to enable the selection of an appropriate hand posture for tool use. When grasp and use hand postures are incongruent, feedback coming by way of the inferior frontal gyrus/premotor cortex modulates activation levels to inhibit the selection of the task-inappropriate grasp hand posture in favor of the task-appropriate tool use hand posture (Buxbaum, 2017).

Outside of the tool domain, few lesion-symptom mapping studies have investigated the neuroanatomic regions critical for selecting an appropriate motor response in the face of conflict. Abnormal interference in the flanker task has been reported to result from lesions to the right premotor cortex (Rinne et al., 2013), the right inferior parietal lobe (Coulthard et al., 2008), as well as the left (Wessel et al., 2014) and right (Geddes et al., 2014) prefrontal cortex. Damage to the striatum (caudate and putamen) is associated with abnormal response interference in the Simon task (Schmidt et al., 2018), potentially as a result of the role of the striatum in mediating interactions between frontal and motor areas when selecting between competing motor responses (Alexander et al., 1986; Aron et al., 2007; Hoffstaedter et al., 2014; Isherwood et al., 2021; Mestres-Misse et al., 2012). To our knowledge, no prior studies have investigated whether the brain regions critical for performance in these classic response interference tasks overlap with (or differ from) those critical for performance of a tool response selection task, as we do in the present study.

## Materials and Methods

### Participants

Sixty-four chronic left cerebrovascular accident (LCVA) participants (38 males, 26 females; mean age, 58.4 years; standard deviation, 10.3 years; mean education, 14.2 years, standard deviation, 2.8 years) were recruited from the Neuro-Cognitive Research Registry at Moss Rehabilitation Research Institute (Schwartz et al., 2005). Participants completed a battery of cognitive and action-based neuropsychological tests and an MRI scan. The presence of limb apraxia was determined based on performance 2 standard deviations below the mean of 16 age-matched healthy controls on a test of gesture to the sight of tools (mean accuracy = 0.94%, standard deviation = 0.06, cut-off score = 0.82%; see Metzgar et al., 2022; Watson & Buxbaum, 2015). Thirty-three of the 64 participants were identified as exhibiting apraxia. We assessed performance in a (1) Grasp-use conflict tool use pantomiming task (hereafter, Grasp-use conflict task), a (2) Flanker task, and a (3) Simon task. The majority of LCVA participants were right-hand dominant (1 reported as left-handed; 1 reported as ambidextrous), and all had suffered a single left hemisphere stroke at least 3 months prior to testing. Participants were excluded if they had a history of psychosis, drug or alcohol abuse, co-morbid neurological disorder, or severe language comprehension deficits established with the Western Aphasia Battery (Kertesz, 1982). The magnitude of the difference between the high and low conflict condition (hereafter, the interference effect) was inspected for outliers with a Shapiro-Wilks test using the ‘check_normality’ function in R. Of the 64 LCVA participants, participant 47’s interference effect in the Simon task was identified as an outlier, and their data were removed from all further analyses. See Table 1 for demographic information and apraxia status of participants, Figure 1 for the cortical and subcortical lesion distribution of each LCVA participant, and Figure 2 for a lesion overlap map.

**Table 1.** Behavioral performance and demographic information for each LCVA participant. Mean accuracy in the flanker task and Simon task are provided in parentheses. *Abbreviations*. ACC, accuracy; RT, response time.

| LCVA<br>Participant | Grasp-Use Conflict Task |  | Flanker Task |  | Simon Task |  | Lesion<br>Volume<br>(1mm<sup>3</sup>) | Education | Gender | Age |
| --- | --- | --- | --- | --- | --- | --- | --- | --- | --- | --- |
|  | High Conflict<br>ACC | Low Conflict<br>ACC | High Conflict<br>RT | Low Conflict<br>RT | High Conflict<br>RT | Low Conflict<br>RT |  |  |  |  |
| 1 | 0.89 | 0.86 | 835 (1.00) | 815 (1.00) | 1055 (0.98) | 918 (1.00) | 80532 | 16 | F | 55 |
| 2 | 0.79 | 0.90 | 719 (1.00) | 686 (1.00) | 1083 (1.00) | 1067 (1.00) | 62530 | 13 | F | 61 |
| 3* | 0.74 | 0.90 | 1158 (1.00) | 928 (1.00) | 916 (0.92) | 716 (0.98) | 5509 | 16 | M | 64 |
| 4 | 0.90 | 1.00 | 725 (0.98) | 609 (1.00) | 681 (0.97) | 633 (1.00) | 166393 | 13 | M | 60 |
| 5* | 0.80 | 0.85 | 1037 (1.00) | 1004 (0.98) | 1235 (1.00) | 1033 (1.00) | 85734 | 14 | M | 61 |
| 6 | 0.94 | 0.95 | 846 (1.00) | 788 (1.00) | 770 (1.00) | 679 (1.00) | 41502 | 14 | F | 58 |
| 7* | 0.47 | 0.60 | 1008 (1.00) | 931 (1.00) | 1182 (1.00) | 1094 (1.00) | 18528 | 12 | F | 81 |
| 8* | 0.65 | 1.00 | 918 (1.00) | 926 (1.00) | 1532 (0.98) | 1461 (0.98) | 47566 | 12 | M | 62 |
| 9 | 0.78 | 1.00 | 842 (0.95) | 842 (0.98) | 809 (0.92) | 684 (0.95) | 64284 | 12 | M | 65 |
| 10* | 0.60 | 0.81 | 1003 (0.95) | 885 (0.98) | 764 (0.98) | 733 (1.00) | 109181 | 16 | F | 68 |
| 11* | 0.68 | 0.68 | 941 (0.98) | 694 (1.00) | 841 (0.95) | 702 (1.00) | 72797 | 18 | F | 40 |
| 12 | 0.85 | 1.00 | 690 (1.00) | 584 (1.00) | 749 (1.00) | 653 (1.00) | 14382 | 12 | F | 68 |
| 13* | 0.70 | 0.86 | 749 (1.00) | 720 (1.00) | 1067 (0.85) | 1099 (0.97) | 50459 | 14 | M | 55 |
| 14* | 0.68 | 0.95 | 881 (0.95) | 810 (1.00) | 823 (0.97) | 807 (1.00) | 89427 | 12 | F | 62 |
| 15 | 0.95 | 0.95 | 752 (1.00) | 663 (1.00) | 841 (0.98) | 808 (1.00) | 56373 | 18 | M | 68 |
| 16* | 0.37 | 0.35 | 1147 (0.95) | 995 (0.95) | 1150 (1.00) | 1110 (1.00) | 71905 | 18 | M | 73 |
| 17 | 0.80 | 0.90 | 1106 (0.98) | 970 (0.98) | 1044 (1.00) | 1061 (1.00) | 126448 | 12 | M | 54 |
| 18 | 0.80 | 0.90 | 881 (0.93) | 862 (0.95) | 961 (0.97) | 869 (1.00) | 44493 | 18 | F | 47 |
| 19 | 0.70 | 1.00 | 655 (0.95) | 587 (1.00) | 692 (0.93) | 648 (0.97) | 55990 | 16 | F | 49 |
| 20 | 0.75 | 1.00 | 840 (0.95) | 751 (0.95) | 850 (0.95) | 776 (1.00) | 48459 | 13 | F | 56 |
| 21* | 0.75 | 0.52 | 724 (0.98) | 612 (1.00) | 917 (0.78) | 734 (0.82) | 13834 | 14 | M | 59 |
| 22* | 0.50 | 0.62 | 625 (0.9) | 562 (0.95) | 728 (0.95) | 645 (0.98) | 5953 | 16 | M | 65 |
| 23 | 0.85 | 0.95 | 731 (1.00) | 763 (0.98) | 589 (0.98) | 568 (1.00) | 181199 | 16 | F | 40 |
| 24 | 0.89 | 1.00 | 573 (0.98) | 538 (1.00) | 789 (0.98) | 717 (1.00) | 96147 | 16 | M | 49 |
| 25 | 0.95 | 0.95 | 729 (1.00) | 656 (1.00) | 861 (0.98) | 868 (0.98) | 337 | 16 | M | 56 |
| 26* | 0.53 | 0.90 | 760 (0.93) | 692 (0.95) | 902 (0.79) | 915 (0.73) | 56281 | 9 | F | 69 |
| 27 | 0.95 | 0.95 | 718 (1.00) | 646 (1.00) | 1089 (1.00) | 1004 (1.00) | 4095 | 18 | M | 64 |
| 28 | 0.95 | 1.00 | 561 (1.00) | 534 (1.00) | 646 (0.95) | 634 (1.00) | 29052 | 12 | M | 61 |
| 29 | 0.84 | 0.89 | 748 (1.00) | 649 (0.98) | 770 (1.00) | 780 (0.98) | 102522 | 16 | F | 39 |
| 30 | 0.85 | 0.95 | 991 (0.95) | 774 (1.00) | 1247 (1.00) | 1073 (1.00) | 14660 | 18 | M | 54 |
| 31 | 0.80 | 1.00 | 947 (0.88) | 929 (0.98) | 980 (0.96) | 894 (1.00) | 26504 | 12 | F | 51 |
| 32 | 0.83 | 0.90 | 1006 (0.95) | 932 (1.00) | 1285 (0.86) | 1189 (0.91) | 87120 | 18 | M | 76 |
| 33 | 0.90 | 0.95 | 533 (1.00) | 496 (1.00) | 656 (0.97) | 547 (1.00) | 189767 | 16 | M | 59 |
| 34 | 0.95 | 0.90 | 615 (1.00) | 582 (0.98) | 785 (1.00) | 723 (0.98) | 39337 | 13 | F | 67 |
| 35* | 0.53 | 1.00 | 659 (1.00) | 610 (1.00) | 690 (0.98) | 645 (1.00) | 32105 | 12 | M | 63 |
| 36 | 0.90 | 0.95 | 1016 (0.98) | 1009 (1.00) | 851 (0.97) | 733 (0.98) | 50394 | 9 | M | 45 |
| 37* | 0.55 | 0.95 | 1019 (0.95) | 927 (0.93) | 1430 (0.98) | 1186 (1.00) | 99980 | 13 | M | 51 |
| 38* | 0.35 | 0.70 | 901 (1.00) | 933 (1.00) | 1070 (1.00) | 1045 (1.00) | 115118 | 16 | F | 59 |
| 39 | 0.90 | 0.85 | 764 (1.00) | 610 (1.00) | 709 (1.00) | 671 (1.00) | 48305 | 12 | F | 36 |
| 40* | 0.60 | 0.90 | 760 (0.98) | 677 (1.00) | 907 (0.98) | 826 (0.98) | 258736 | 18 | M | 73 |
| 41* | 0.55 | 0.74 | 820 (1.00) | 717 (1.00) | 808 (0.97) | 761 (0.98) | 69778 | 12 | M | 57 |
| 42* | 0.60 | 0.85 | 753 (1.00) | 711 (1.00) | 847 (0.98) | 893 (1.00) | 76301 | 18 | M | 71 |
| 43 | 0.74 | 0.95 | 831 (1.00) | 771 (1.00) | 1191 (1.00) | 1159 (1.00) | 51780 | 21 | M | 72 |
| 44 | 0.95 | 1.00 | 562 (0.95) | 561 (0.98) | 793 (0.97) | 708 (0.97) | 17706 | 12 | M | 51 |
| 45* | 0.47 | 0.53 | 1010 (1.00) | 863 (0.95) | 1065 (1.00) | 1084 (1.00) | 5350 | 14 | M | 49 |
| 46* | 0.40 | 0.39 | 1004 (0.93) | 1070 (0.93) | 1391 (1.00) | 1286 (0.93) | 225021 | 16 | M | 64 |
| 47*# | 0.47 | 0.76 | 709 (0.98) | 603 (1.00) | 1465 (0.87) | 962 (1.00) | 179606 | 15 | M | 63 |
| 48 | 1.00 | 1.00 | 557 (0.98) | 499 (1.00) | 836 (1.00) | 838 (1.00) | 20105 | 21 | M | 57 |
| 49* | 0.74 | 0.84 | 653 (1.00) | 637 (0.98) | 1041 (1.00) | 1035 (1.00) | 64793 | 14 | M | 67 |
| 50* | 0.47 | 0.79 | 916 (0.98) | 885 (0.95) | 1613 (0.87) | 1447 (0.95) | 61198 | 12 | F | 72 |
| 51* | 0.42 | 0.90 | 972 (1.00) | 829 (1.00) | 1256 (0.7) | 1199 (0.73) | 136576 | 12 | F | 46 |
| 52* | 0.70 | 0.95 | 917 (0.90) | 920 (0.95) | 1153 (0.95) | 1364 (0.88) | 85091 | 12 | M | 49 |
| 53* | 0.37 | 0.50 | 954 (1.00) | 802 (1.00) | 1644 (0.67) | 1741 (0.73) | 147237 | 12 | F | 30 |
| 54* | 0.60 | 0.79 | 580 (0.95) | 533 (1.00) | 826 (1.00) | 814 (1.00) | 136565 | 16 | M | 55 |
| 55* | 0.60 | 0.80 | 626 (1.00) | 608 (1.00) | 953 (0.97) | 917 (1.00) | 200079 | 12 | M | 62 |
| 56 | 0.95 | 1.00 | 944 (0.90) | 835 (0.98) | 1263 (0.97) | 1233 (1.00) | 190665 | 13 | M | 54 |
| 57* | 0.53 | 0.74 | 1095 (0.92) | 823 (0.88) | 887 (0.95) | 768 (0.98) | 121243 | 12 | F | 47 |
| 58 | 0.83 | 0.95 | 835 (0.98) | 874 (0.93) | 1199 (0.97) | 1176 (1.00) | 16977 | 9 | M | 51 |
| 59* | 0.55 | 0.85 | 1000 (0.85) | 895 (0.93) | 931 (0.98) | 941 (1.00) | 32684 | 13 | F | 64 |
| 60 | 0.79 | 1.00 | 708 (0.98) | 594 (1.00) | 1014 (0.98) | 1018 (0.98) | 30414 | 12 | F | 68 |
| 61* | 0.39 | 0.79 | 845 (1.00) | 781 (1.00) | 1130 (1.00) | 1120 (1.00) | 202614 | 15 | F | 59 |
| 62* | 0.50 | 0.60 | 768 (1.00) | 668 (1.00) | 837 (1.00) | 754 (1.00) | 32003 | 18 | M | 71 |
| 63 | 0.79 | 0.95 | 655 (0.88) | 601 (1.00) | 820 (0.95) | 760 (1.00) | 27004 | 12 | M | 49 |
| 64* | 0.75 | 0.89 | 785 (0.98) | 733 (0.93) | 1491 (0.97) | 1377 (1.00) | 1869 | 12 | F | 67 |

\* Tool use gesturing performance was two standard deviations below the mean of age-matched controls (averaging across high and low conflict conditions).

**Figure 1.**
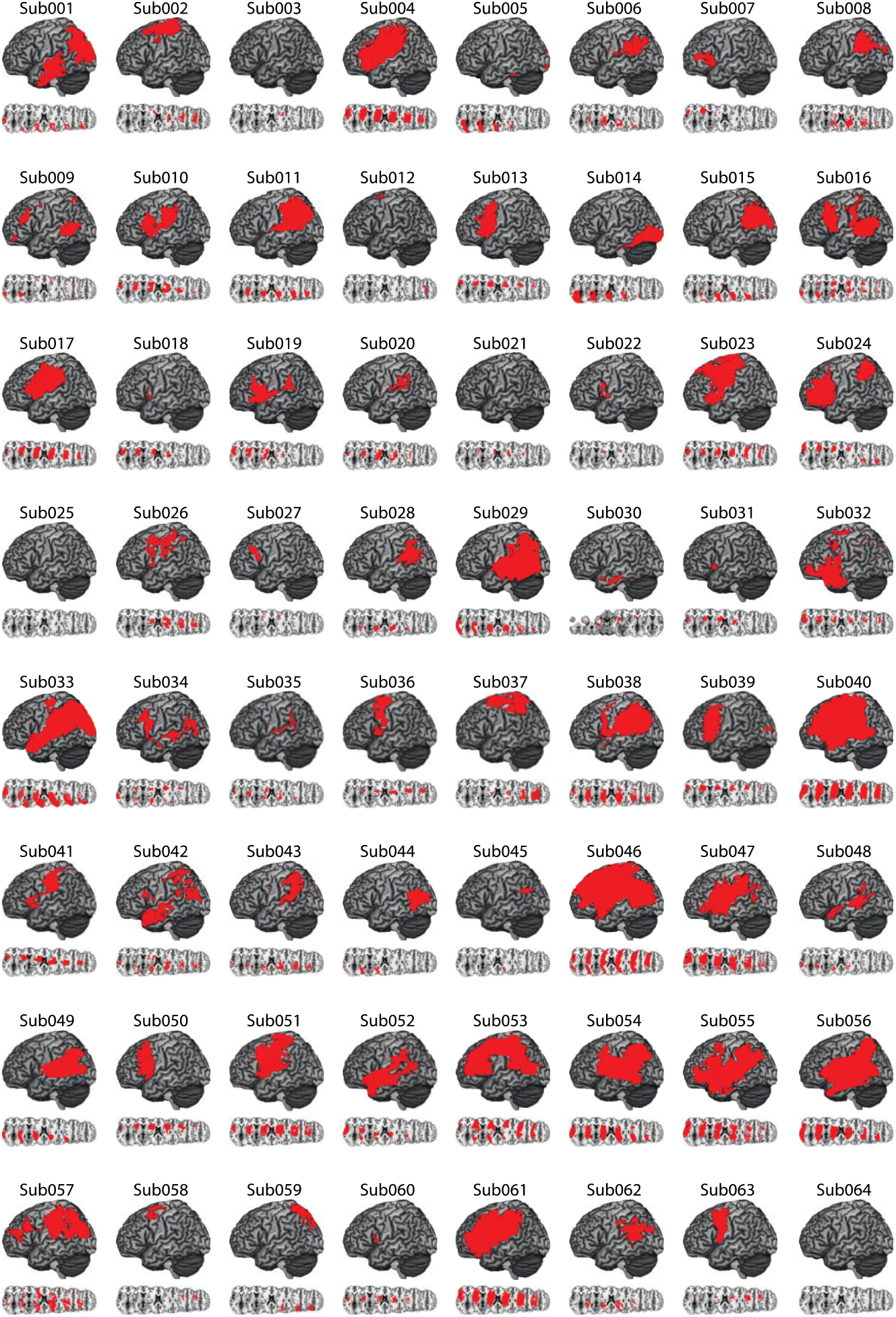
Distribution of Cortical and Subcortical Damage in each LCVA Participant. A depiction of the 64 LCVA participants’ lesions are presented on the ch2bet template. Lesions are projected on the cortical surface but include subcortical voxels using a 12-voxel search depth. Axial slices begin at the MNI coordinate [0,0,0] and increase in 10-mm increments superiorly.

**Figure 2.**
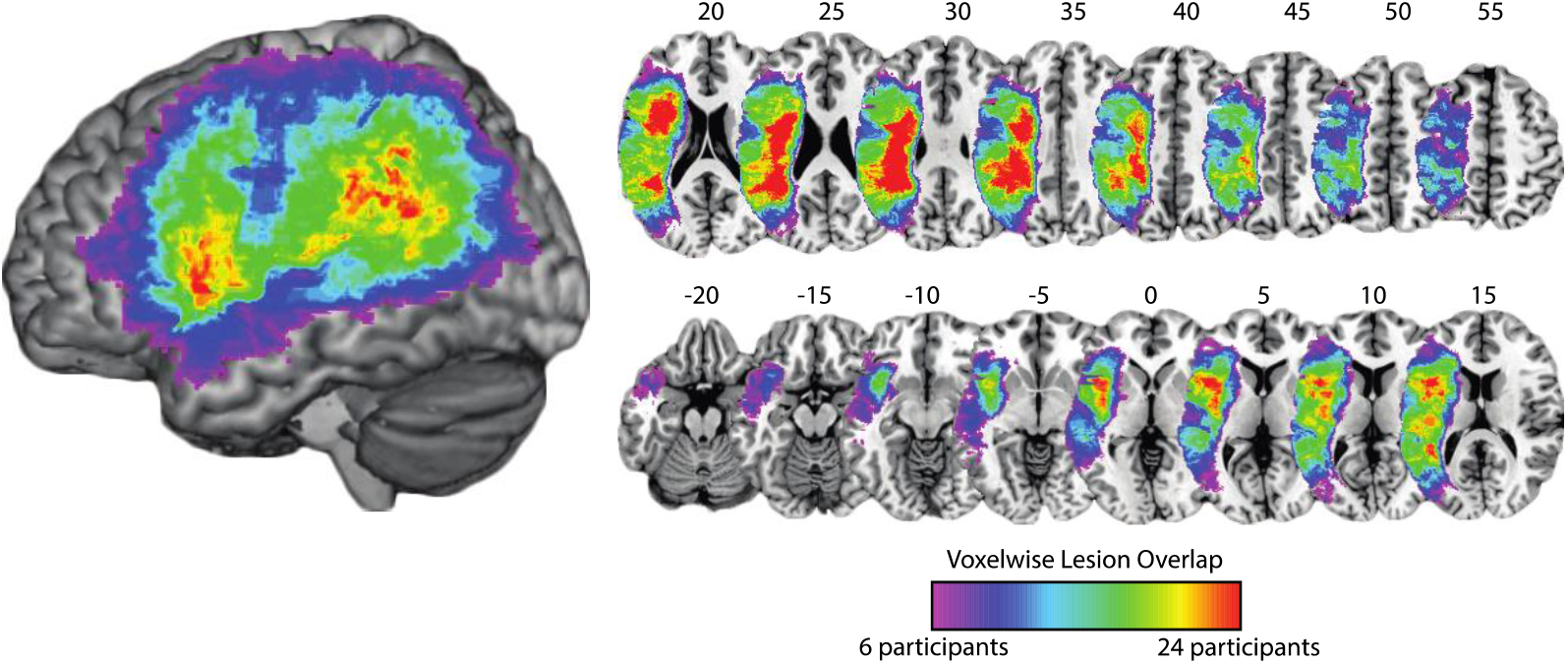
Voxelwise overlap map among LCVA participants. Lesion overlap among the 63 LCVA participants is projected on the surface representation of the ch2bet brain (left) and in volume space (right). MNI Z coordinates are listed for each axial slice.

### Neuropsychological Testing

#### Grasp-Use Conflict Task

The grasp-use conflict task included 40 photographs of manipulable objects (tools) taken from the BOSS database (Brodeur et al., 2010). Tools included items with distinct use actions, including construction tools (e.g., wrench), household articles (e.g., teapot), office supplies (e.g., scissors), and bathroom items (e.g., razor). Half of the items had conflicting hand position/actions for use versus grasp (i.e., high conflict items, e.g., corkscrew), while half of the items had a single hand position/action for both use and grasp (i.e., low conflict items, e.g., hammer). High and low conflict tools differed significantly in conflict rating (*t*(38) = 10.00, *p* < 0.001; Hedges’ G (HG) effect size, 3.10), but were equivalent in affordance strength (*t*(38)=1.50, *p* = 0.14; HG, 0.45), name agreement (*t*(38) = 1.13, *p* = 0.26; HG, 0.35), and familiarity (*t*(38) = 1.50, *p* = 0.13; HG, 0.45; for details, see Watson & Buxbaum, 2015; Garcea et al., 2019). Each trial of the grasp-use conflict task began with the presentation of a 400-by-400 pixel color photograph of a tool on a computer monitor. Participants were asked to “show how you would use the tool as if you were holding and using it” with the left (non-motorically-impaired) hand. Four practice trials with feedback (using items different than in the task itself) were given at the start of the task. As per Rothi et al. (1991), if a participant gestured the action as if their hand was the tool (body-part-as-object error), they were reminded to “show how you would use the tool as if you were actually holding it in your hand”. The first of these errors was corrected and the participant was permitted a second try.

#### Coding of Tool Use Gesturing Data

The grasp-use conflict task was video-recorded with a digital camera and actions were scored offline by two trained, reliable coders (Cohen’s Kappa score = 94%) who also demonstrated inter-rater reliability with previous coders in the Buxbaum lab (Cohen’s Kappa > 85%; see e.g., Buxbaum et al., 2005b). Both tasks were coded using a portion of the detailed praxis scoring guidelines used in our group’s previous work (Buxbaum et al., 2000; Buxbaum, Kyle, et al., 2005; Watson & Buxbaum, 2015). First, each gesture was given credit for semantic content unless a participant performed a recognizable gesture appropriate for a semantically related tool. Only gestures that were given credit for semantic content were scored on the spatiotemporal hand action dimension. A spatiotemporal hand action error was assigned if the shape or movement of the hand and/or wrist was flagrantly incorrect, or if the hand or finger was used as part of the tool (i.e., body-part-as-object error; Buxbaum et al., 2005b; Watson & Buxbaum, 2015). We calculated the mean spatiotemporal hand posture accuracy for high and low conflict tool use actions for each LCVA participant (see Table 1 and Figure 3A).

**Figure 3.**
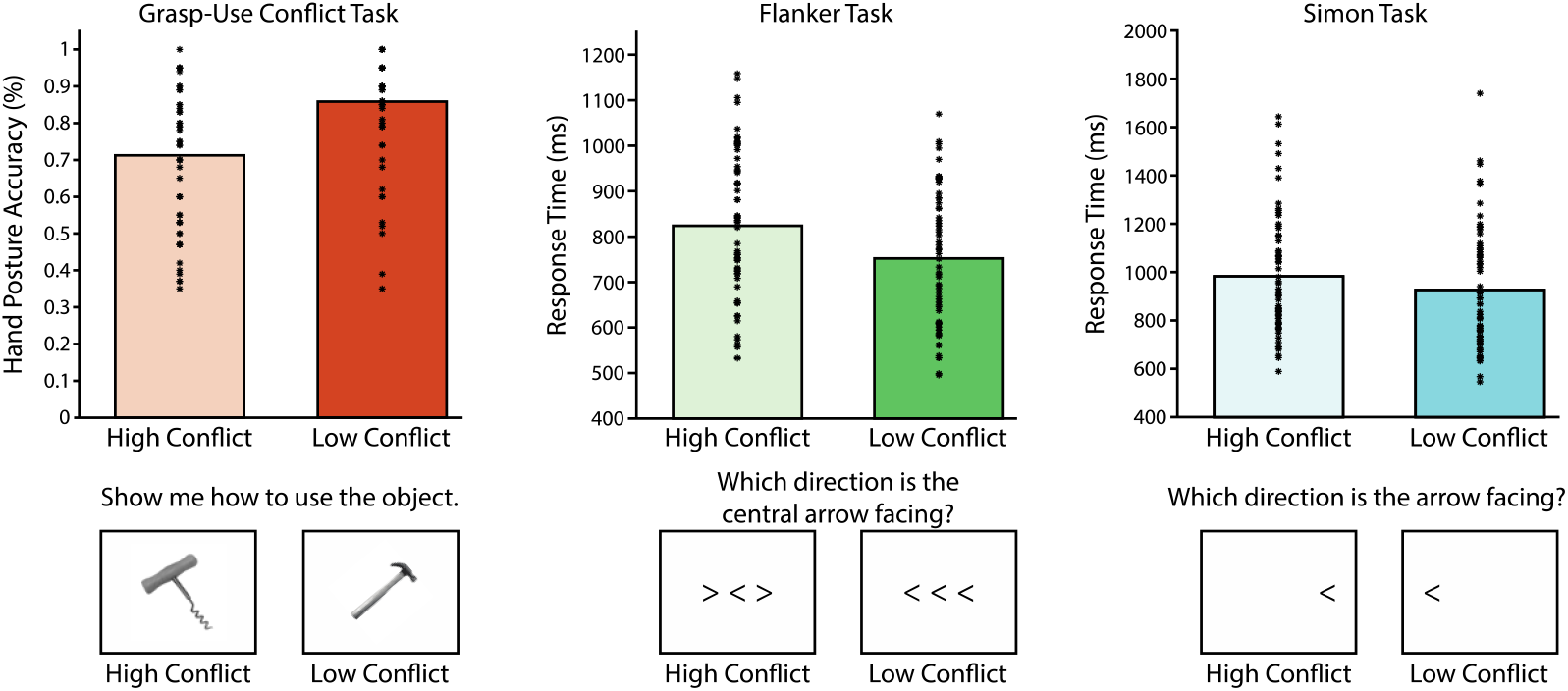
Accuracy and RT measures of response selection across tasks. Mean accuracy (grasp-use conflict task) and RTs (flanker task; Simon task) for high and low conflict conditions are represented as bar plots. Each asterisk represents an individual participant’s mean accuracy or RT across trials.

Using linear regression, we controlled for shared variance between performance in the high and low conflict conditions and entered residual high conflict scores as the dependent variable in a support vector regression lesion-symptom mapping analysis. Negative-going residual values indicate that participants were differentially less accurate to gesture the use of high conflict tools in relation performance when gesturing the use of low conflict tools.

#### Flanker Task

A publicly available version^1^ of the flanker task was used to investigate stimulus-response selection. There were six cells in the design: Arrow Direction (two levels; left, right) and Flanking Arrow Conflict (three levels; high conflict, low conflict, neutral). On each trial, a right- or left-facing arrow was presented in the center of the screen and participants were directed to use the index or middle finger to indicate whether the arrow was facing left or right, respectively. Participants used their left index and middle finger to ensure that responses were generated with the unaffected hand/fingers. On low conflict trials, two flanking arrows were facing the same direction as the central arrow (e.g., < < <), whereas on high conflict trials the flanking arrows were facing the opposite direction (e.g., > < >). Neutral trials required participants to make a left/right decision of a central arrow presented in isolation. All participants completed 12 trials of training to familiarize themselves with the task prior to the experiment. The experiment proper contained 120 trials (20 trials per cell of the design) presented randomly.

Response time (RT) data on accurate trials were trimmed to remove trials greater than 2.5 standard deviations above or below the mean separately for high conflict, low conflict, and neutral conditions (197 trials (2.60%) were removed. We then collapsed across arrow direction to obtain a mean RT and accuracy for high and low conflict trials (see Table 1 and Figure 3B; the neutral condition is not germane to the current investigation and is not analyzed here). Mean RTs were then log-transformed. Linear regression was used to control for shared variance between high and low conflict conditions, and residual high conflict RTs were entered as the dependent variable in a support vector regression lesion-symptom mapping analysis. Positive-going residual values indicate that participants were differentially slower to categorize arrow direction in the high conflict condition in relation to performance in the low conflict condition.

#### Simon Task

A Simon task was developed using Psyscope stimulus presentation software. There were 6 cells in the design: Visual Field Location (two levels; left, right) and Visual Field Conflict (three levels; high conflict, low conflict, neutral). On each trial, a right- or left-facing arrow was presented in the right visual field, left visual field, or in the center of the visual field. Participants were instructed to use the left index or middle finger to indicate whether the arrow was facing left or right, respectively^2^. Participants used their left index and middle finger to ensure that RTs were generated with the unaffected hand/fingers. On low conflict trials the arrow direction was congruent with the visual field in which it was presented in (e.g., a right-facing arrow presented in the right visual field), whereas on high conflict trials the arrow direction was facing the opposite direction (e.g., a right-facing arrow presented in the left visual field). Neutral trials required participants to make a left or right decision of a centrally presented arrow. Participants completed 18 trials of training to familiarize themselves with the task prior to the experiment proper. The experiment proper contained 180 trials presented randomly within a block of 60 trials (10 trials per cell of the design per 60 trials presented randomly).

RT data on accurate trials were trimmed to remove trials greater than 2.5 standard deviations above or below the mean separately for high conflict, low conflict, and neutral conditions (306 trials (2.69%) were removed. We then collapsed across arrow direction to obtain a mean RT and accuracy for high and low conflict trials (see Table 1 and Figure 3C; the neutral condition is not germane to the current investigation and is not analyzed here). Mean RTs were then log-transformed. Linear regression was used to control for shared variance between high and low conflict conditions, and residual high conflict RTs were entered as the dependent variable in a support vector regression lesion-symptom mapping analysis. Positive-going residual values indicate that participants were differentially slower to categorize arrow direction in the high conflict condition in relation to their performance in the low conflict condition.

### Neuroimaging Acquisition

#### Acquisition of Anatomic Scans

MRI scans included whole-brain T1-weighted MR images collected on a 3T (Siemens Trio, Erlangen, Germany; repetition time = 1620 ms, echo time = 3.87 milliseconds, field of view = 192 × 256 mm, 1 × 1 × 1 mm voxels) or a 1.5T (Siemens Sonata, repetition time = 3,000 milliseconds, echo time = 3.54 milliseconds, field of view = 24 cm, 1.25 × 1.25 × 1.25 mm voxels) scanner, using an eight-channel or sixty-four channel head coil. Lesions were manually segmented on each LCVA participant’s high-resolution T1-weighted structural images. Lesioned voxels, consisting of both grey and white matter, were assigned a value of 1 and preserved voxels were assigned a value of 0. Binarized lesion masks were then registered to a standard template (Montreal Neurological Institute “Colin27”) using a symmetric diffeomorphic registration algorithm (https://picsl.upenn.edu/software/ants/) (Avants et al., 2008).

Volumes were first registered to an intermediate template comprised of healthy brain images acquired on the same scanner. Volumes were then mapped onto the “Colin27” template to complete the transformation into standard space. To ensure accuracy during the transformation process, lesion maps were subsequently inspected by a neurologist (H.B. Coslett), who was naïve to the behavioral data of the study. For increased accuracy, the pitch of the template was rotated to approximate the slice plane of each LCVA participant’s scan. This method has been demonstrated to achieve high intra- and inter-rater reliability (Schnur et al., 2009). See Table 1 for lesion size quantification.

### Support Vector Regression Lesion Symptom Mapping (SVR-LSM) Analyses

SVR-LSM was performed in MATLAB 2017B using a toolbox developed by DeMarco and Turkeltaub (2018). SVR-LSM is a multivariate technique that uses machine learning to determine the association between lesioned voxels and behavior when considering the lesion status of all voxels submitted to the analysis. It overcomes several limitations of univariate VLSM, including inflated false positives from correlated neighboring voxels (Pustina et al., 2018), type II error due to correction for multiple comparisons (Bennett et al., 2009), and uneven statistical power due to biased lesion frequency as a function of vascular anatomy (Mah et al., 2014 Sperber & Karnath, 2017). SVR-LSM has been shown to be superior to VLSM when multiple brain areas are involved in a single behavior (Herbet et al., 2015; Mah et al., 2014; Mirman, Zhang, et al., 2015; Sperber et al., 2019; Zhang et al., 2014).

Only voxels lesioned in at least 10% of participants (6 participants) were included. We controlled for variability in lesion volume using the ‘Direct Total Lesion Volume Control’ corrective method (DeMarco & Turkeltaub, 2018). Five-fold cross-validation was implemented, in which 80% of the participants’ lesions and behavioral data were used to train a classifier, and the remaining 20% of participants’ lesions and behavioral data were used to test the classifier. This procedure was iterated 5 times to ensure that each unique subset of participant data was independently used for training and testing, and the resulting 5 maps of feature weights were averaged together to derive a final map of voxelwise beta values. Voxelwise statistical significance was then determined using a Monte Carlo style permutation analysis in which the behavioral data were randomly assigned to a lesion map, and the same procedure as described above was iterated 10,000 times. Voxelwise z-scores were then computed for the true data in relation to the mean and standard deviation of voxelwise null distributions; the resulting z-score map was set to a threshold of z < -1.65 (*p* < .05, one-tailed) to determine chance-level likelihood of a lesion-symptom relation. We then further restricted the resulting map by eliminating any clusters with fewer than 500 contiguous voxels (Garcea et al., 2020; Lacey et al., 2017; Skipper-Kallal et al., 2017). The Anatomical Automatic Labeling (AAL) atlas was used to assess the location of significant voxels (Tzourio-Mazoyer et al., 2002).

## Data availability statement

In compliance with the guidelines of the Institutional Review Board of Einstein Healthcare Network, all participants gave informed consent and were compensated for travel expenses and participation. The informed consents obtained did not include permission to make data publicly available. Accordingly, the conditions of our ethical approval do not permit anonymized raw data to be publicly archived. To obtain access to the data, individuals should contact the corresponding author. Requests for data are assessed and approved by the Institutional Review Board of Einstein Healthcare Network. The data summarized in Table 1 and the R scripts used for analysis can be found at https://github.com/frankgarcea/GarceaBuxbaum_ActionSelectionProject.git.

## Results

### Behavioral Interference Effects

There were robust interference effects (less accurate or slower performance in the high conflict condition compared to the low conflict condition) in the grasp-use conflict task (*t*(62) = 8.37, *p* < .001), flanker task (*t*(62) = 8.54, *p* < .001), and Simon task (*t*(62) = 6.18, *p* < .001; see Figure 3 and Table 2).

**Table 2.**
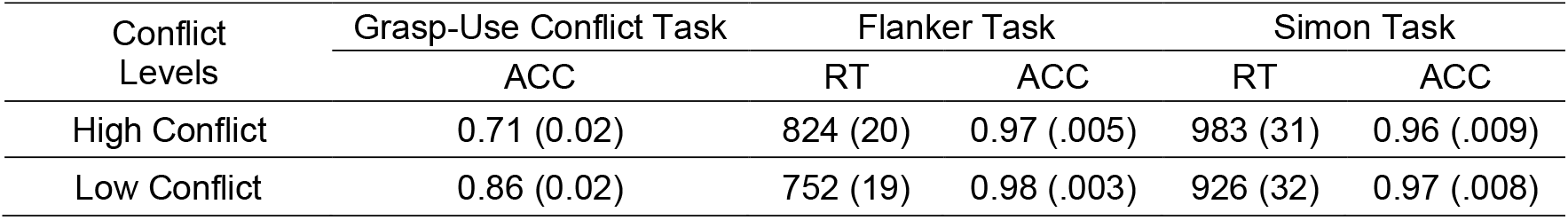
Mean group-level accuracy or RTs as a function of high and low conflict conditions across tasks. Standard error of the mean computed across participants is listed in parentheses. Abbreviations: ACC, accuracy; RT, response time.

| Conflict Levels | Grasp-Use Conflict Task | Flanker Task |  | Simon Task |  |
| --- | --- | --- | --- | --- | --- |
|  | ACC | RT | ACC | RT | ACC |
| High Conflict | 0.71 (0.02) | 824 (20) | 0.97 (.005) | 983 (31) | 0.96 (.009) |
| Low Conflict | 0.86 (0.02) | 752 (19) | 0.98 (.003) | 926 (32) | 0.97 (.008) |

### Correlations Within Tasks

There was a significant correlation between hand posture performance accuracy when gesturing the use of high and low conflict tools (*r*(61) = 0.68, *p* < .001). Similarly, there were significant correlations between high and low conflict RTs on the flanker task (*r*(61) = 0.91, *p* < .001), as well as on the Simon task (*r*(61) = 0.96, *p* < .001).

### Correlations Between Tasks

Less accurate performance when gesturing the use of high conflict tools was associated with slower RTs in the high conflict condition of the flanker task (r(61) = -0.40, *p* < .005) and Simon task (r(61) = -0.37, *p* < .001). There were also significant correlations between the high conflict conditions of the flanker task and Simon task (r(61) = 0.56, *p* < .001; see Table 3). Qualitatively similar correlations in performance were observed in the low conflict condition across the 3 tasks. However, the correlations between residual high conflict scores (i.e., performance in the high conflict condition controlling for performance in the low conflict condition) were not significant or were only marginally correlated (see Table 3).

**Table 3.** Cross-task correlations between response interference performance.

| Conflict Levels | Grasp-Use Conflict Task and Flanker Task | Grasp-Use Conflict Task and Simon Task | Flanker Task and Simon Task |
| --- | --- | --- | --- |
| High Conflict | $r(61) = -0.37, p < .01$ | $r(61) = -0.39, p < .01$ | $r(61) = 0.53, p < .001$ |
| Low Conflict | $r(61) = -0.34, p < .01$ | $r(61) = -0.27, p < .05$ | $r(61) = 0.52, p < .001$ |
| Residual High Conflict Scores | $r(61) = 0.03, p = 0.84$ | $r(61) = 0.12, p = 0.35$ | $r(61) = 0.20, p = 0.11$ |

### Interference Effects Modulated by Apraxia Status

We then used a mixed effects ANOVA with the between-subject factor of Apraxia Status (two levels; apraxic, non-apraxic) and the within-subject factors of Conflict Level (two levels; high conflict, low conflict) to investigate whether the magnitude of the interference effect was modulated by apraxia status separately for each task. In the Grasp-Use Conflict task, there was a main effect of Group (*F*(1, 61) = 95.20, *p* < .001, η^2^ = 0.61), Conflict Level (*F*(1, 61) = 81.45, *p* < .001, η^2^ = 0.57), and an interaction between the factors (*F*(1, 61) = 11.82, *p* < .001, η^2^ = 0.16). Planned contrasts (*t*-test, two-tailed) demonstrated that the magnitude of the interference effect was significantly larger in individuals with apraxia than in individuals without apraxia (*t*(61) = 4.18, *p* < .001; see Table 4). In the Flanker task, there was a main effect of Group (*F*(1, 61) = 6.48, *p* < .05, η^2^ = 0.10) and Conflict Level (*F*(1, 61) = 89.36, *p* < .001, η^2^ = 0.59), but no interaction between factors (F < 1). Similarly, in the Simon task there was a main effect of Group (*F*(1, 61) = 6.46, *p* < .05, η^2^ = 0.10) and Conflict Level (*F*(1, 61) = 73.17, *p* < .001, η^2^ = 0.55), but no interaction between the factors (*F*(1, 61) = 1.43, *p* = 0.27, η^2^ = 0.02).

**Table 4.**
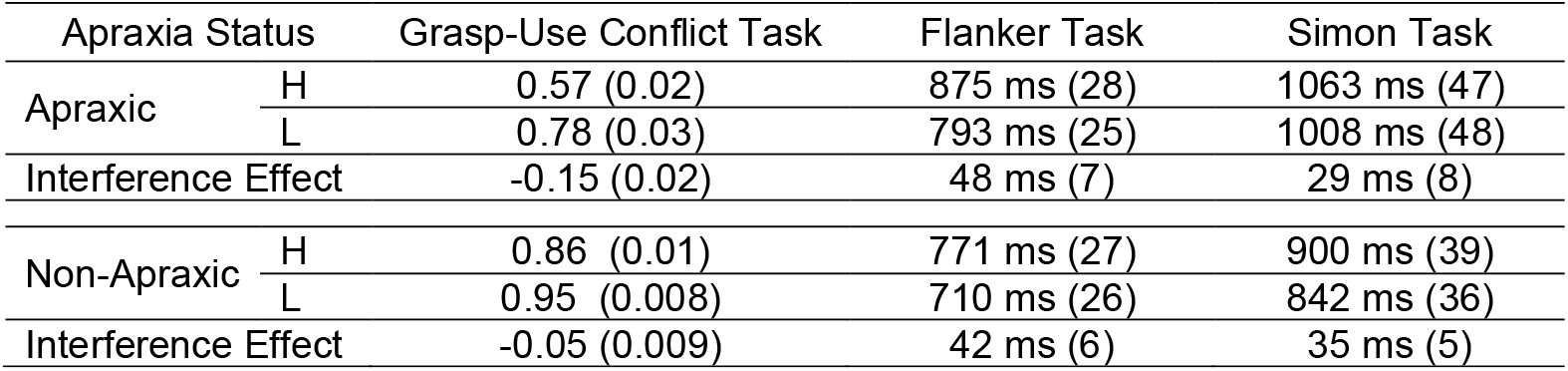
The mean accuracy or response time (in ms) and interference effect are presented separately as a function of apraxia status. Participant-level interference effects were computed by subtracting the mean of the low conflict (L) condition from the mean of the high conflict condition (H), and dividing the difference by the sum of the two scores. Standard error of the mean computed across participants is provided in parentheses.

### Support Vector Regression Lesion Symptom Mapping Results

#### Grasp-Use Conflict Task

The first SVR-LSM analysis tested the hypothesis that disproportionately lower accuracy when gesturing the use of high conflict tools relative to low conflict tools would be associated with lesions to the left inferior parietal lobule, the left inferior frontal gyrus, the left precentral gyrus, and the left insula (Garcea et al., 2019; Watson & Buxbaum, 2015; Watson et al., 2019). Replicating our prior work, we identified lesions to two large clusters associated with differentially poor performance when gesturing the use of high conflict tools (see Figure 4A). The first cluster included the left post-central gyrus, the left posterior superior temporal gyrus, the left inferior parietal lobule (supramarginal gyrus and angular gyrus), the left superior parietal lobule, and the left precuneus. The second cluster included the left pre-central gyrus, the left frontal operculum, the left Rolandic operculum, the left insula, and the left putamen (see Figure 4A and Table 5A). Voxel-level data were significant (z < -1.65, *p* < .05, one-tailed) and survived a 10,000-iteration cluster-level permutation analysis. We identified qualitatively similar results when repeating the analysis with a univariate VLSM approach (see Supplemental Figure 2).

**Table 5.** Peak MNI coordinates identified in the SVR-LSM analysis of the Grasp-Use Conflict Task (A), the Flanker Task (B), and the Simon Task (C). Region labels were derived from the Automated Anatomical Labeling (AAL) atlas.

| **A. Grasp-Use Conflict Task** |  |  |  |  |  |  |
| --- | --- | --- | --- | --- | --- | --- |
| AAL Label | AAL Number | Mean Center of Mass (XYZ) |  |  | Peak Z-Score | Number of Voxels |
| Precentral Gyrus | 1 | -40 | -3 | 28 | 3.12 | 417 |
| Rolandic Operculum | 17 | -41 | -17 | 24 | 3.72 | 972 |
| Insula | 29 | -37 | -18 | 24 | 3.29 | 2553 |
| Postcentral Gyrus | 57 | -22 | -38 | 58 | 3.35 | 5708 |
| Superior Parietal Lobule | 59 | -21 | -50 | 50 | 3.24 | 774 |
| Inferior Parietal Lobule | 61 | -38 | -41 | 41 | 3.72 | 3558 |
| Supramarginal Gyrus | 63 | -48 | -39 | 35 | 2.89 | 933 |
| Putamen | 73 | -26 | 3 | 16 | 2.23 | 64 |
| Heschel's Gyrus | 79 | -46 | -16 | 10 | 2.33 | 265 |
| Superior Temporal Gyrus | 81 | -41 | -9 | -9 | 2.69 | 554 |
| **B. Flanker Task** |  |  |  |  |  |  |
| AAL Label | AAL Number | Mean Center of Mass (XYZ) |  |  | Peak Z-Score | Number of Voxels |
| Insula | 29 | -34 | 3 | 13 | 2.35 | 62 |
| Postcentral Gyrus | 57 | -39 | -32 | 46 | 3.35 | 4386 |
| Inferior Parietal Lobule | 61 | -54 | -24 | 45 | 3.72 | 2683 |
| Supramarginal Gyrus | 63 | -59 | -24 | 42 | 3.72 | 1186 |
| Caudate Nucleus | 71 | -18 | 9 | 24 | 2.70 | 411 |
| Putamen | 73 | -24 | 2 | 16 | 3.24 | 723 |
| **C. Simon Task** |  |  |  |  |  |  |
| AAL Label | AAL Number | Mean Center of Mass (XYZ) |  |  | Peak Z-Score | Number of Voxels |
| Caudate Nucleus | 71 | -18 | -8 | 26 | 3.35 | 425 |
| Putamen | 73 | -21 | 1 | 14 | 2.47 | 488 |
| Pallidum | 75 | -25 | -4 | 2 | 2.03 | 174 |

**Figure 4.**
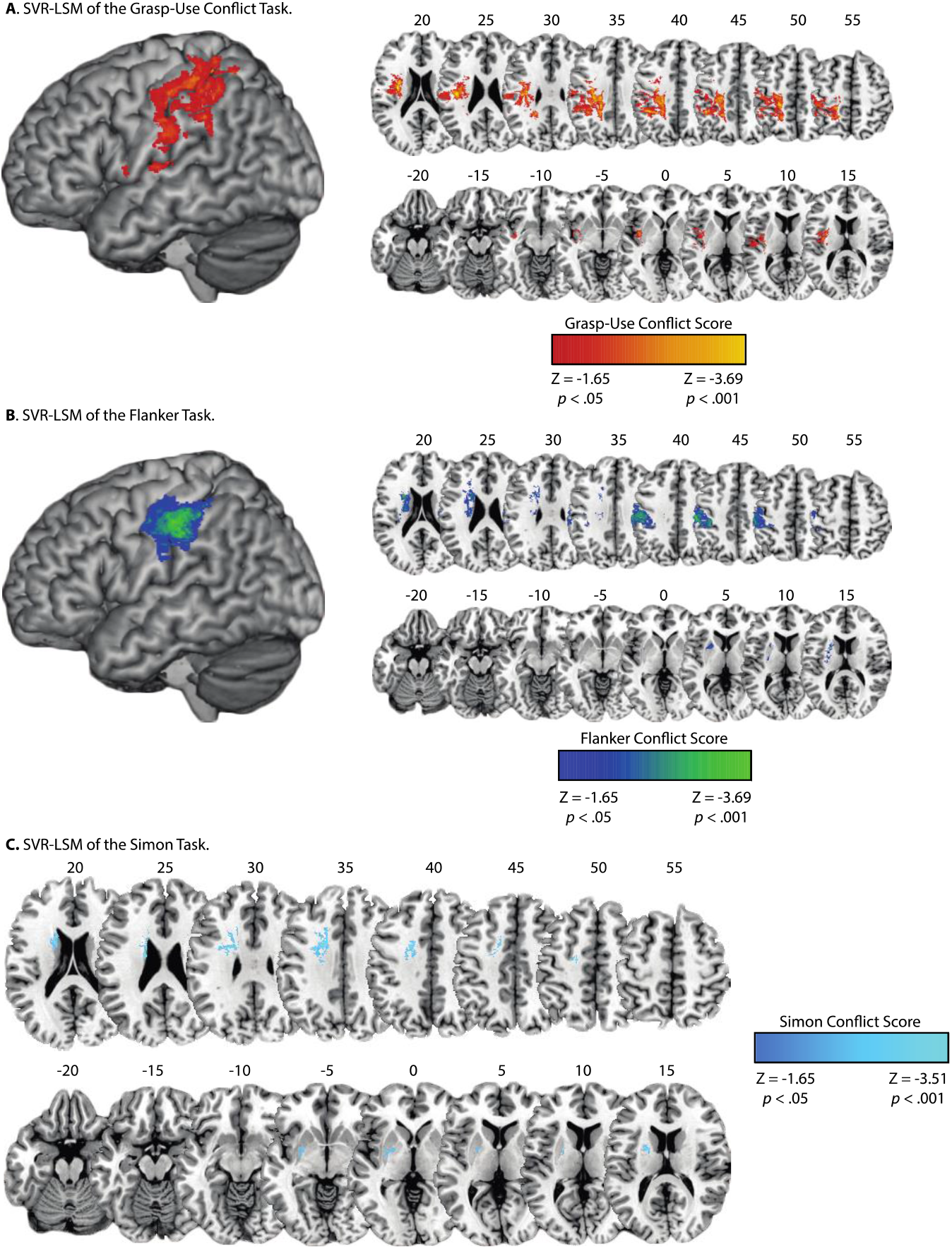
SVR-LSM of action selection in tool and non-tool tasks. Surface-based (left) and volumetric (right; MNI Z coordinate listed above axial slices) renderings of the voxels associated with **(A)** Reduced performance (lower accuracy when gesturing the use of high conflict tools controlling for performance when gesturing the use of low conflict tools) in the grasp-use conflict task, **(B)** Increased RTs (slower RTs in the high conflict condition controlling for RTs in the low conflict condition) in the flanker task, and **(C)** Increased RTs (slower RTs in the high conflict condition controlling for RTs in the low conflict condition) in the Simon task.

#### Flanker Task

The SVR-LSM analysis of the flanker task tested the hypothesis that disproportionately slower RTs in the high conflict condition, controlling for variance in the low conflict condition, would be associated with lesions to the left inferior and superior parietal lobule and subcortical regions including the caudate and putamen (Rinne et al., 2013). We identified lesions to two large clusters that were associated with differentially slower RTs in the high conflict condition (see Figure 4B). The first cluster included the left post-central gyrus, the left inferior parietal lobule (supramarginal gyrus), and the left superior parietal lobule. The second cluster included the left insula, caudate, putamen, and pallidum (see Figure 4B and Table 5B). Voxel-level data were significant (z < -1.65, *p* < .05, one-tailed) and formed clusters of at least 500 contiguous voxels. We identified qualitatively similar results when repeating the analysis using a univariate VLSM approach (see Supplemental Figure 3).

#### Simon Task

The SVR-LSM analysis of the Simon task tested the hypothesis that disproportionately slower RTs in the high conflict condition, controlling for variance in the low conflict condition, would be associated with lesions to subcortical regions including areas of the left caudate, putamen, and insula (Schmidt et al., 2018). We identified lesions to a cluster of areas that was associated with differentially slower RTs in the high conflict condition (see Figure 4C). The cluster included the white matter undercutting the middle frontal gyrus, and the caudate, putamen, and pallidum (see Figure 4C and Table 5C). Voxel-level data were significant (z < -1.65, *p* < .05, one-tailed) and formed clusters of at least 500 contiguous voxels. We identified qualitatively similar results when repeating the analysis using a univariate VLSM approach (Supplemental Figure 4).

### Overlapping Lesion Sites Associated with Abnormal Response Selection

To quantify the overlap between tasks, we computed a voxelwise overlap ratio of the SVR-LSM results using AAL defined regions-of-interest (ROI; c.f. Glascher et al., 2012). The overlap ratio was calculated by computing the number of overlapping voxels within an ROI between tasks relative to the sum of all significant voxels within an ROI between tasks [overlap ratio = (sum of overlapping voxels in maps A and B in an ROI) / (sum of all significant voxels in maps A and B in an ROI)]. We interpret the overlap between tasks if the ratio exceeds 1% in a region. Three ROIs were identified in the overlap analysis between the grasp-use conflict task and the flanker task (see Figure 5A): (1) The left postcentral gyrus (11.5%), (2) The left inferior parietal lobule (6.2%), and (3) The left supramarginal gyrus (2%).^3^ Two ROIs were identified in the overlap analysis between the flanker task and Simon task (see Figure 5B): (1) The left caudate nucleus (26%), and (2) the left putamen (3%). No ROIs were identified in the overlap analysis of the grasp-use conflict task and the Simon task.

**Figure 5.**
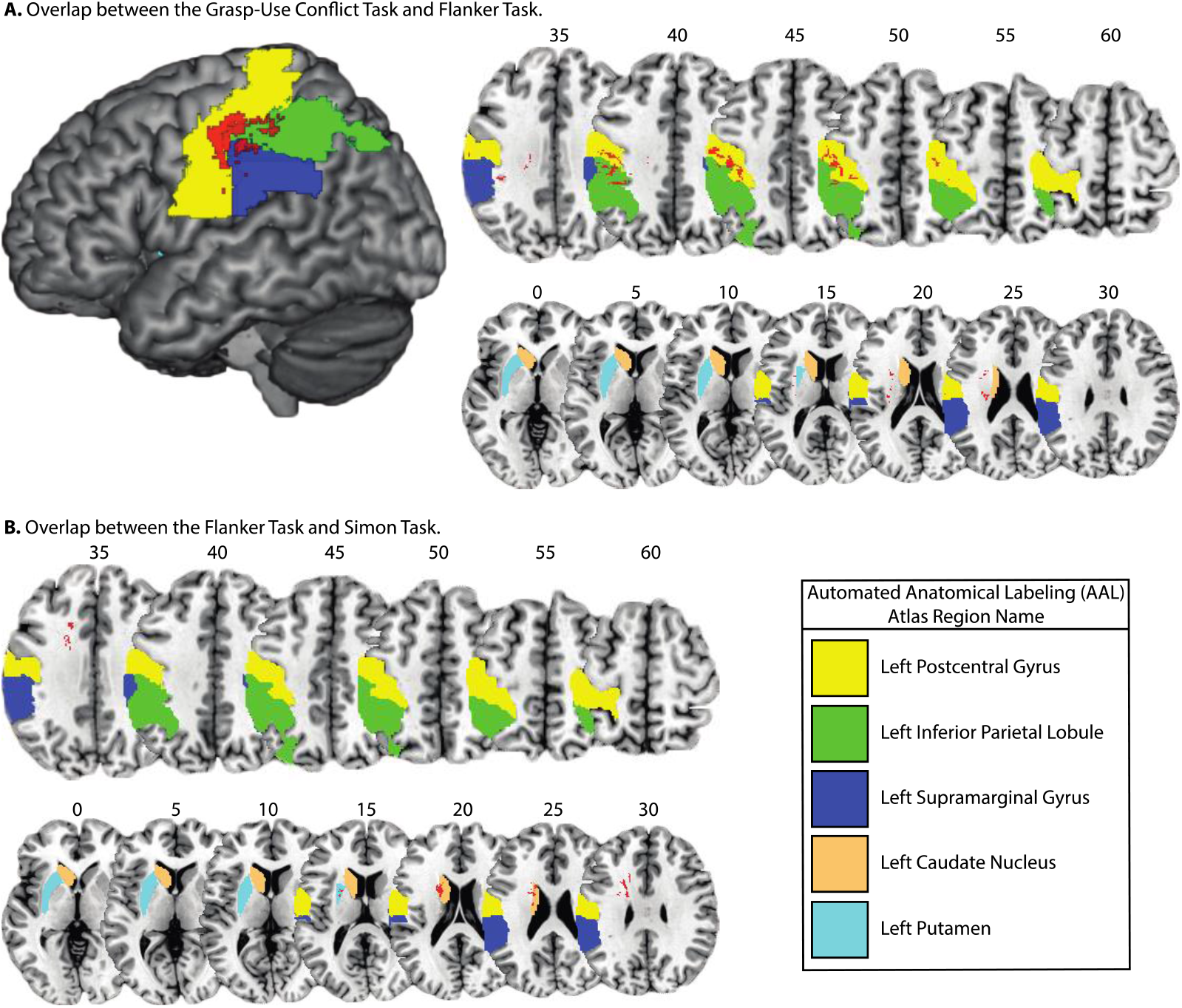
Overlap between voxels associated with response interference in tool and non-tool tasks. (A) Surface-based (left) and volumetric (right; MNI Z coordinate listed above axial slices) renderings of voxelwise overlap between the grasp-use conflict task and the flanker task. There was overlap in the left postcentral gyrus, the left inferior parietal lobule (supramarginal gyrus), and in subcortical voxels adjacent to the caudate nucleus and putamen. (B) Voxelwise overlap between the flanker task and Simon task was identified in the caudate nucleus and putamen, and the white matter undercutting the middle frontal gyrus.

## General Discussion

On a moment-to-moment basis, interacting with tools in the environment requires selection of an appropriate action from multiple possible actions. In the case of tools with incongruent hand postures for grasping-to-move and functional use (high conflict tools, e.g., screwdriver), individuals with apraxia after left hemisphere stroke are abnormally error-prone. This abnormal pattern has been associated with lesions to the supramarginal gyrus, insula, and inferior frontal gyrus/ventral premotor cortex (Watson & Buxbaum, 2015). This study was the first to assess the relation between grasp-use conflict effects and response interference on non-tool tasks in LCVA participants, 33 of whom exhibited apraxia.

We observed that performance on high-conflict conditions was correlated across tool, flanker, and Simon tasks, yet correlations between residual high conflict scores (i.e., high conflict controlling for variance in low-conflict conditions) were not significant (e.g., see Table 3; see also Glascher et al., 2012). Moreover, the magnitude of the interference effect was modulated by apraxia status in the Grasp-Use Conflict task but not in the flanker or Simon task. These data are consistent with prior studies reporting absent or weak correlations in the magnitude of the flanker and Simon response interference effects (e.g., Hedge et al., 2021; Wager et al., 2005), suggesting that interference effects may not rest on domain-general cognitive control mechanisms such as capacity to inhibit irrelevant responses. One explanation for the absence of correlations between interference effects in the present study is that our tasks incorporate different loci of conflict between stimulus and response dimensions. In the Simon task, conflict arises between different responses tied to stimulus location and stimulus arrow direction, whereas in the flanker task, conflicting responses are associated with the arrow directions of two stimuli (i.e., central and flanking arrows). In the tool use gesturing task, conflicting responses are associated with visual shape-based object “affordances” and functional manipulation knowledge. The behavioral data are thus consistent with the suggestion that correlated interference effects are not observed when dimensional overlap between tasks is low (Whitehead et al., 2019; see also Kornblum, 1992; Kornblum, 1994; Kornblum et al., 1990).

Consistent with prior studies from our lab (e.g., Watson & Buxbaum, 2015; Garcea et al., 2019; Watson et al., 2019), the present SVR-LSM data indicate that lesions to the left inferior parietal lobule, the left ventral premotor cortex, and the left insula are associated with relatively reduced tool use pantomiming accuracy when LCVA participants gestured the use of high conflict tools relative to low conflict tools. We investigated whether these lesion sites would be associated with abnormal response conflict in the Simon and flanker tasks, both classic non-tool response conflict paradigms. Differentially slower RTs in the high conflict condition of the Erikson flanker task were associated with lesions to the left inferior and superior parietal lobule, the left postcentral gyrus, the left insula, and the left caudate and putamen. In the Simon task, differentially slower RTs in the high conflict condition were associated with lesions to the left caudate and putamen, and white matter undercutting the left precentral gyrus and left middle frontal gyrus. All SVR-LSM results were replicated using a traditional univariate VLSM approach. Overlap analyses identified the left inferior parietal lobule, the left postcentral gyrus, and the white matter lateral to the left caudate and left putamen as common lesion sites associated with reduced response selection performance across grasp-use conflict and flanker tasks. Lesions involving the left caudate and left putamen were associated with reduced response selection across the flanker and Simon tasks. These results are discussed in the context of theoretical models of praxis function and response selection in the human brain.

### Response Selection in the Service of Skilled Action Production

Prior functional MRI (fMRI) studies have identified a left-lateralized network of brain regions that supports skilled action production, referred to as the Tool Use Network (Almeida et al., 2013; Garcea & Buxbaum, 2019; Lesourd et al., 2021). The regions include the posterior middle temporal gyrus (pMTG), supramarginal gyrus (SMG), and ventral premotor cortex (PMv) bordering on the inferior frontal (IFG). The left pMTG is argued to support the representation and retrieval of tool use knowledge (Lingnau & Downing, 2015; Martin, 2007), including representations of the appearance of the hand postures and body movements that are associated with familiar tools (e.g., that a spoon is transported along a particular trajectory for eating; Buxbaum et al., 2014 Tarhan et al., 2015). The left SMG supports the production of gestures, including meaningless gestures and tool use actions (Johnson-Frey, 2004; Orban & Caruana, 2014). The PMv is involved the planning and implementation of compound or multi-step tool use actions (Chao & Martin, 2000; Garcea & Buxbaum, 2019; Grafton et al., 1997) and in the sequencing of action (De Renzi et al., 1983; Harrington & Haaland, 1992). Whereas impaired tool use pantomiming in apraxia has been associated with lesions to each of these areas (Dressing et al., 2018; Garcea et al., 2013; Goldenberg, 2009; Martin et al., 2016; Osiurak et al., 2021; Randerath et al., 2009; Schmidt et al., 2022; Weiss et al., 2001), susceptibility to grasp-on-use interference has been linked to lesions involving the left SMG, PMv/IFG, insula, and underlying white matter, including the superior longitudinal fasciculus (Garcea et al., 2019; Watson & Buxbaum, 2015). In a study of LCVA participants, the degree of grasp-on-use interference was associated with reduced resting state functional connectivity between the left SMG and the left PMv (Watson et al., 2019). These prior data, in tandem with the current findings, are consistent with a neurocognitive model of tool use in which the left SMG serves to buffer candidate hand posture representations during the response selection process. Modulatory input that comes by way of the left PMv biases processing such that task-irrelevant hand postures are inhibited, and task-relevant hand postures are selected for tool use (Buxbaum, 2017; Garcea et al., 2019).

Our neurocognitive model dovetails with the theoretical framework of Rounis & Humphreys (2015), which leveraged an affordance competition framework (Cisek & Kalaska, 2010) to postulate that the left inferior parietal lobule is a key substrate for resolving competition between different tool actions. The investigators argued that apraxia can arise from the selection of a task-irrelevant tool use action that shares similar visuomotor features with a task-relevant action. For example, hammer and butcher knife are two tools that share similar visual affordances (e.g., a long handle that is grasped to use the object) and action kinematics (e.g., swung along a similar trajectory) despite being used in different contexts to satisfy different behavioral goals. The affordance competition framework stipulates that top-down modulatory input is required to inhibit a task-irrelevant action (e.g., chopping meat with a butcher knife) when intending to select a similar tool use action (e.g., pounding in a nail with a hammer). Work from our laboratory has shown that individuals with apraxia after posterior temporal or parietal lesions exhibit a reduction in the degree to which tools with similar use actions compete for selection (Garcea et al., 2019; Lee et al., 2013). Moreover, the frequency of overt gestural errors is correlated with the degree of slowing of activation of these representations (Lee et al., 2013). These data suggest that weakness of activation of tool use representations affects the likelihood that they will be rapidly selected, and thus reduces their ability to guide overt behavior (Buxbaum & Kalenine, 2021; Garcea et al., 2019; Watson & Buxbaum, 2015).

Across all tasks, LCVA participants with abnormal response selection had lesions involving the putamen or both caudate and putamen (see Table 5). The computational role of the caudate and putamen in praxis is unclear considering that apraxia is rarely associated with frank lesions to these structures (Leiguarda & Marsden, 2000; Pramstaller & Marsden, 1996). Damage to these areas in the right hemisphere has been associated with abnormal selection of grip strength force in the presence of distracting visual stimuli (Rinne et al., 2018). Using the flanker task, Wylie et al. (2009) demonstrated that there were larger interference effects in participants with Parkinson’s disease relative to neurotypical controls. This finding suggests that degradation of the striatum can lead to heightened response conflict when inhibiting the selection of an irrelevant action (see also Verbruggen & Logan, 2008; Wylie et al., 2010). Interestingly, lesions to the left caudate and putamen have been associated with larger interference effects in the Simon task in stroke survivors (Schmidt et al., 2018). Moreover, a prior study using an fMRI version of the Simon paradigm reported increased BOLD contrast in the caudate nucleus bilaterally when neurotypical participants made incongruent judgments (Schmidt et al., 2020). Our SVR-LSM and univariate LSM results are consistent with these findings and suggest that caudate and putamen damage can give rise to less efficient selection of task-relevant representations across tool and non-tool domains (for review, see Isherwood et al., 2021).

An important avenue for future brain mapping research is to determine whether lesions to the white matter undercutting the left PMv/IFG can lead to reduced grasp-use response selection. For example, lesions to a white matter bottleneck undercutting the IFG are associated with semantic memory retrieval deficits in absence of IFG damage (Mirman, Chen, et al., 2015; see also Griffis et al., 2017). In praxis, it has been argued that a “ventral” pathway supporting tool use pantomime connects the left inferior frontal gyrus (BA 45) with the left inferior parietal lobule via fibers passing through the extreme capsule (Hoeren et al., 2014; Vry et al., 2015). These fibers are in close anatomical proximity to the caudate and putamen. It therefore remains a possibility that impaired selection of tool use responses may arise from disconnection of inferior parietal areas from frontal areas via damage to fibers passing through the extreme capsule. Case-series based studies (Metzgar et al., 2022) are needed to investigate grasp-use conflict effects in LCVA participants with lesions isolated to the caudate and putamen in the absence of damage to the extreme capsule. This approach will permit fine-grained analyses of the lesion sites and structural and functional connectivity among brain regions supporting efficient and flexible cognitive control of skilled action in the human brain.

## Conclusion

In summary, we did not observe significant correlations in the response interference effects across tool and non-tool tasks. Although there was partial lesion-symptom mapping overlap between tasks, we did not identify a common lesion site associated with the magnitude of the response interference effect across all 3 tasks. These results, in the context of prior research, suggest that the grasp-use interference effect is a specific hallmark of apraxia that is not reducible to a domain-general difficulty with response selection, but rather arises from deficits in the activation and selection of tool use representations specifically.

## Acknowledgements

We thank Cortney Howard, Harrison Stoll, and Veronica Kreter for coding participants’ gestures, Austin Wild, Olu Faseyitan and Branch Coslett for help with lesion drawing and segmentation, and Christine Watson for help organizing the behavioral and lesion data. Preparation of this manuscript was supported by a Moss Rehabilitation Research Institute/University of Pennsylvania postdoctoral training fellowship (NIH 5T32HD071844-05), and by NIH grant R01 NS099061 to L.J.B. F.E.G. is supported by the University of Rochester CTSA awards NIH UL TR002001 and NIH KL2 TR001999 from the National Center for Advancing Translational Sciences.

## Author Information

The authors declare no competing financial interests. Correspondence should be addressed to Frank E. Garcea.

## SUPPLEMENTAL ONLINE MATERIALS

### Analysis of Simon Interference Effects for each Stimulus Presentation Program

Eight of the 63 LCVA participants took part in a version of the Simon task using E-Prime software. The mean interference effect (high conflict RT – low conflict RT) of those eight participants was marginally larger (mean = 94 ms; red vertical bar in Supplemental Figure 1) than the mean interference effect (mean = 52 ms; standard deviation = 24 ms) of a 10,000-iteration random selection of 8 of the 55 participants who took part in a PsyScope version of the task (see Supplemental Figure 1). For each iteration, 8 randomly selected PsyScope participants’ interference effects were computed; after iterating the analysis 10,000 times we derived a distribution from which to z-score the mean E-Prime interference effect. The z-score associated with a 94 ms effect was marginally significant (z = 1.76, p = 0.08).

We then performed a chi square analysis to determine if the number of participants who exhibited a interference effect larger than the mean of the 10,000-iteration distribution was different across the PsyScope and E-Prime groups. All participants were grouped by the stimulus presentation program (0 = PsyScope; 1 = E-Prime), and interference effects were binarized (1 if greater than 52 ms; 0 if less than 52 ms); these vectors were entered into the ‘crosstab’ function in Matlab. The resulting chi square analysis did not reach significance (χ = 0.65, *p* = 0.43), indicating that although the mean magnitude of the 8 E-Prime participants’ interference effect was marginally greater than the mean of 8 randomly selected PsyScope participants, the proportion of participants whose interference effects were above or below a mean of 52 ms was not different as a function of stimulus presentation program (see Supplemental Table 1).

**Supplemental Table 1.** Proportion of participants who exhibited a interference effect above or below the mean of the 10,000-iteration analysis as a function of stimulus presentation program.

|  | Interference Effect < 52 ms | Interference Effect > 52 ms | Proportion > 52 ms |
| --- | --- | --- | --- |
| E-Prime | 3 | 5 | 5/8 (63%) |
| PsyScope | 29 | 26 | 26/55 (47%) |

**Supplemental Figure 1.**
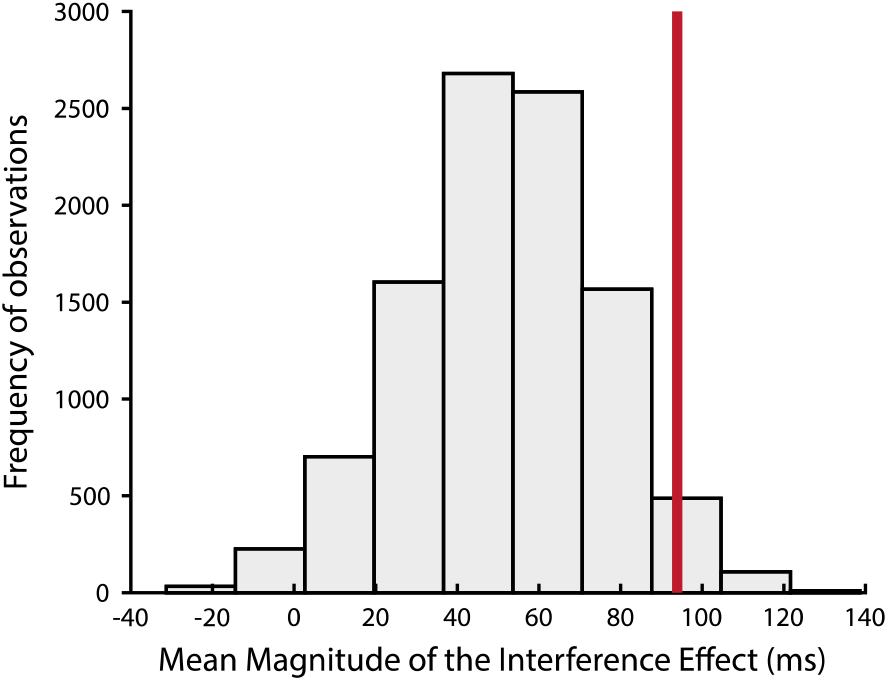
Inspection of Simon Interference Effects as a Function of Stimulus Presentation Program. A 10,000-iteration procedure was carried out in which we randomly selected 8 LCVA participants who took part in a PsyScope version of the Simon task. The distribution represents the mean interference effect (high conflict RT – low conflict RT) for each iteration. The mean interference effect of the 8 E-Prime participants (94 ms) is represented by the red vertical line.

**Supplemental Figure 2.**
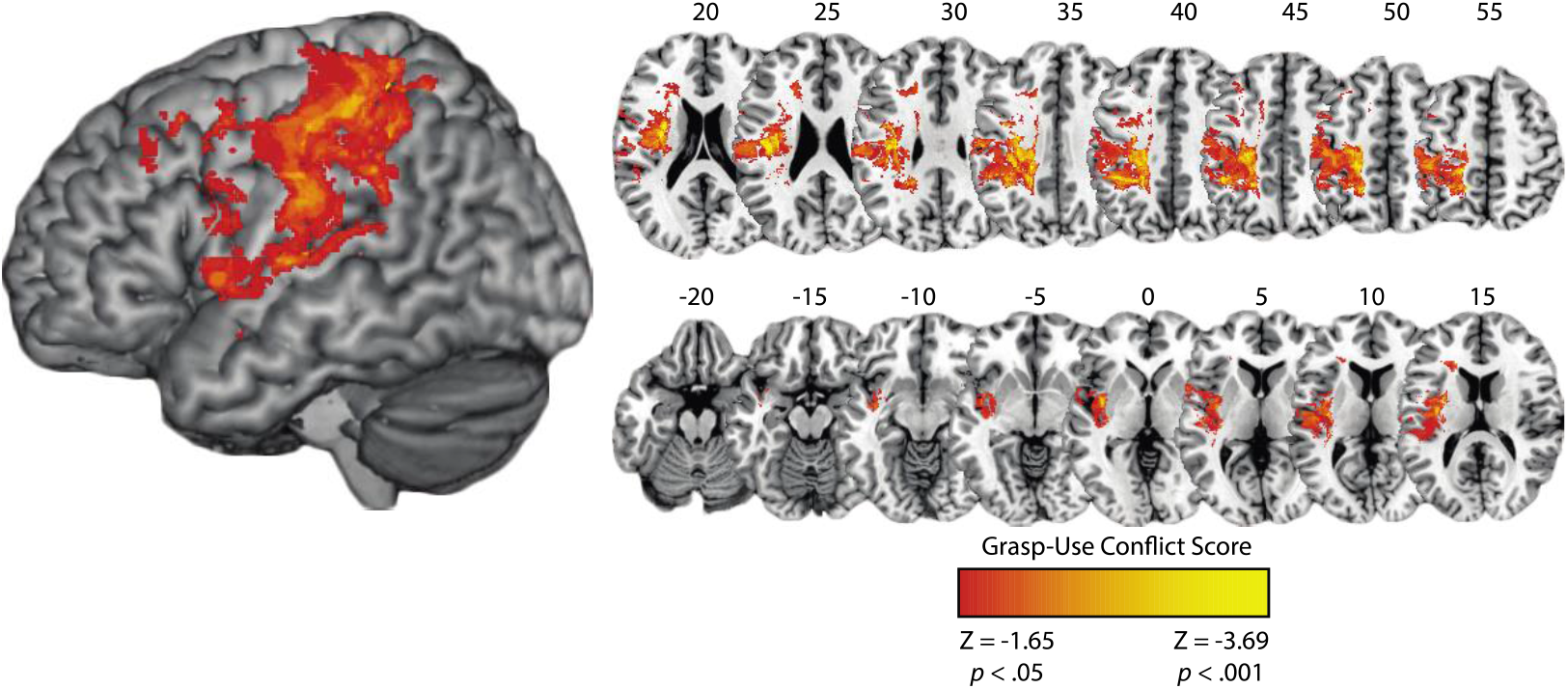
VLSM of the Grasp-Use Conflict Task. **(A)** Cortical (left) and subcortical (right; MNI Z coordinate listed above axial slices) renderings of the VLSM result identifies comparable lesion sites as was observed in Figure 4A.

We identified lesions to three large clusters that were associated with differentially poorer tool use gesturing performance with high conflict tools (see Supplemental Figure 2). The first cluster included lesion sites involving the post-central gyrus, the posterior superior temporal gyrus, the posterior middle temporal gyrus, the inferior parietal lobule (supramarginal gyrus and angular gyrus), superior parietal lobule, the precuneus, and paracentral lobule. The second cluster included lesions sites involving the pre-central gyrus, pars opercularis, pars triangularis, the Rolandic operculum, the insula, and subcortical structures including the caudate nucleus and putamen. The third cluster included lesion sites involving the supplementary motor area, tissue of the middle cingulum, the superior frontal gyrus, and middle frontal gyrus (see Supplemental Table 2). Note that these voxel-level data survive voxel-level multiple comparisons correction and a 10,000-iteration cluster-level permutation analysis.

**Supplemental Table 2.** Peak MNI coordinates identified in the VLSM analysis of increased grasp-use conflict in tool use gesturing. Region labels were derived from the Automated Anatomical Labeling (AAL) atlas.

| AAL Label | AAL Number | Mean Center of Mass (XYZ) |  |  | Peak Z-Score | Number of Voxels |
| --- | --- | --- | --- | --- | --- | --- |
| Precentral Gyrus | 1 | -41 | -2 | 28 | -3.72 | 3048 |
| Superior Frontal Gyrus | 3 | -13 | 20 | 49 | -3.16 | 57 |
| Middle Frontal Gyrus | 7 | -32 | 23 | 46 | -2.63 | 543 |
| Pars Opercularis | 11 | -38 | 9 | 20 | -2.52 | 254 |
| Rolandic Operculum | 17 | -41 | -17 | 24 | -3.72 | 2466 |
| Insula | 29 | -34 | -11 | 16 | -3.72 | 4731 |
| Postcentral Gyrus | 57 | -21 | -38 | 58 | -3.72 | 12786 |
| Superior Parietal Lobule | 59 | -19 | -46 | 62 | -3.72 | 1119 |
| Inferior Parietal Lobule | 61 | -38 | -41 | 40 | -3.72 | 5852 |
| Supramarginal Gyrus | 63 | -47 | -38 | 36 | -3.54 | 2044 |
| Putamen | 73 | -26 | 3 | 16 | -2.86 | 156 |
| Heschl's Gyrus | 79 | -43 | -18 | 8 | -3.06 | 1067 |
| Superior Temporal Gyrus | 81 | -42 | -13 | -2 | -3.19 | 2950 |

**Supplemental Figure 3.**
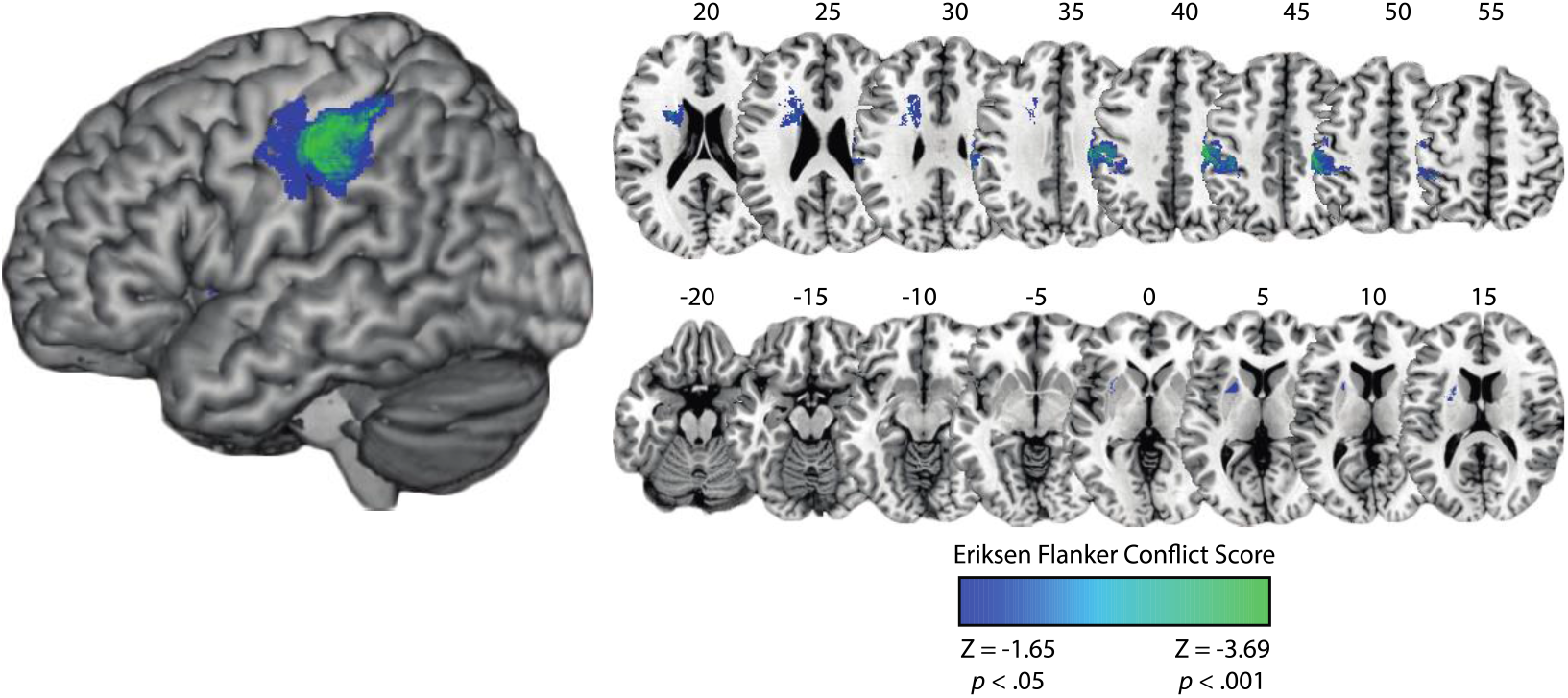
VLSM of the Eriksen Flanker Task. **(A)** Cortical (left) and subcortical (right; MNI Z coordinate listed above axial slices) renderings of the VLSM result identifies comparable lesion sites as was observed in Figure 4B.

We identified lesions to two large clusters that were associated with differentially slower response times in the high conflict condition. The first cluster included lesion sites involving the pre-central and post-central gyri, the inferior parietal lobule (supramarginal gyrus), and the superior parietal lobule. The second cluster included lesions sites involving the insula, the caudate nucleus, putamen, and pallidum (see Supplemental Table 3). Voxel-level data survive voxelwise correction at z < -1.65, *p* < .05, and form of a cluster of at least 500 contiguous voxels.

**Supplemental Table 3.** Peak MNI coordinates identified in the VLSM analysis of the Eriksen flanker task. Region labels were derived from the Automated Anatomical Labeling (AAL) atlas

| AAL Label | AAL Number | Mean Center of Mass (XYZ) |  |  | Peak Z-Score | Number of Voxels |
| --- | --- | --- | --- | --- | --- | --- |
| Postcentral Gyrus | 57 | -13 | 20 | 49 | -3.72 | 2765 |
| Inferior Parietal Lobule | 61 | -38 | 9 | 20 | -3.54 | 3121 |
| Supramarginal Gyrus | 63 | -36 | 23 | 19 | -3.54 | 1280 |
| Caudate Nucleus | 71 | -41 | -17 | 24 | -2.48 | 393 |
| Putamen | 73 | -13 | 10 | 49 | -2.61 | 588 |

**Supplemental Figure 4.**
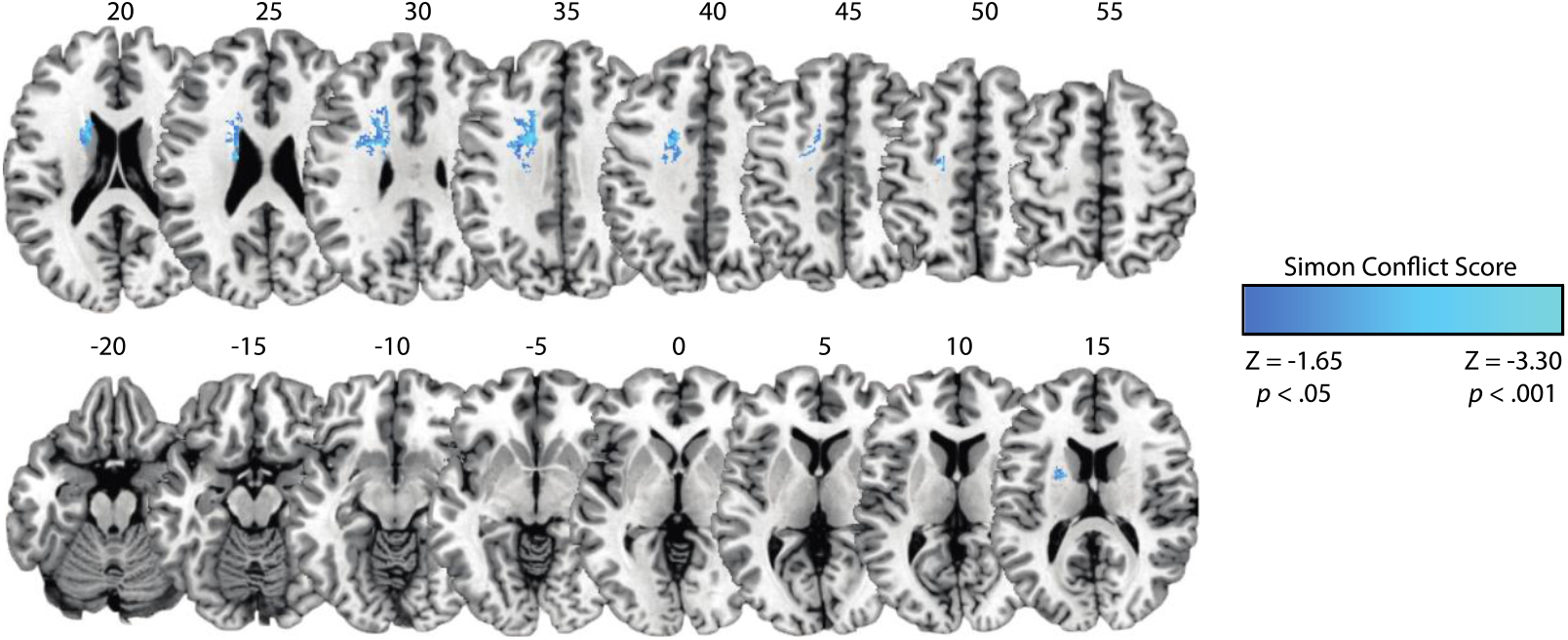
VLSM of the Simon Task. **(A)** A rendering of the VLSM result identifies comparable lesion sites as was observed in Figure 4C.

Lesions to the caudate and putamen were associated with differentially slower response times in the high conflict condition (see Supplemental Table 4). Voxel-level data survive voxelwise correction at z < -1.65, *p* < .05, and form of a cluster of at least 500 contiguous voxels.

**Supplemental Table 4.** Peak MNI coordinates identified in the VLSM analysis of the Simon task. Region labels were derived from the Automated Anatomical Labeling (AAL) atlas.

| AAL Label | AAL Number | Mean Center of Mass (XYZ) |  |  | Peak Z-Score | Number of Voxels |
| --- | --- | --- | --- | --- | --- | --- |
| Caudate Nucleus | 71 | -18 | -8 | 26 | 3.09 | 493 |
| Putamen | 73 | -21 | 1 | 14 | 2.37 | 146 |

## Footnotes

1 http://pebl.sourceforge.net/wiki/index.php/Flanker_Task

2 A subset of participants took part in a Simon task that used E-Prime software for stimulus presentation. All aspects of stimulus presentation and task instruction was identical. In Supplemental Figure 1 we demonstrate the RT results were comparable across stimulus presentation software programs.

3 The AAL atlas distinguishes between a supramarginal gyrus region and an inferior parietal lobule region. For the purpose of the overlap analysis these regions were analyzed separately.

